# A claim-centred atlas distinguishes biological corroboration from analytical depth in plant stress transcriptomes

**DOI:** 10.64898/2026.10.09.757829

**Authors:** Wenze Ding, Peng Zhang, Yi Ding

**Affiliations:** Plant Breeding Institute, School of Life and Environmental Sciences, University of Sydney, Cobbitty, NSW 2570, Australia

## Abstract

Plant stress transcriptomes are reanalysed far more often than independently replicated, obscuring whether apparent consensus reflects biological corroboration or repeated analysis of the same experiments. Here we present PLEASE (PLant Evidence Atlas for StrEss), a claim-centred atlas that treats the independent experiment, rather than the analysis, as the unit of evidence. PLEASE resolves 88,789 analytical records into 62 dependency-resolved study families. A calibrated core of abiotic and biotic stress programmes persists after dependency collapse. Yet support erodes sharply as claims become more specific. Across stress, organ and cell-lineage contexts, independent-match coverage falls from 1.00 to 0.26 while directional agreement remains near 0.68. Three quarters of sampled contexts contain only one study family. Thus, missing comparable experiments, not pervasive disagreement, is the main barrier to context-specific corroboration. PLEASE separates analytical depth from biological corroboration and pinpoints where independent experiments are most needed.

## Main

Transcriptomes are among the most abundant molecular records of how plants respond to stress. Two decades of profiling, from the AtGenExpress compendium to single-cell and spatial atlases of stressed organs^1–5^, have deposited thousands of stress experiments in public repositories^6,7^, while community atlases and databases now make them broadly searchable^8–10^. These data have informed the identification of conserved stress programmes^11–13^ and the prioritization of targets, tissues and cell types for crop improvement^14,15^. As public transcriptomes are increasingly reused to support synthesis and prioritization, a basic question arises: how many independent experiments actually support a given biological conclusion?

That support is difficult to recover from a literature organized around datasets and publications. Two features obscure it. First, the same biological material can appear under several accessions, support multiple papers and generate hundreds of analytical outputs, while changes in normalization, annotation or gene-set definition test analytical robustness without adding biological replication^16–19^. Second, as conclusions become more specific, a lack of corroboration may simply mean that no independent experiment has sampled the same context. For example, a claim about one programme in one root cell type under one stress^20^ may yield many analytical results when the same experiment is reanalysed, but these still amount to a single biological test. Corroboration requires replication in a separate experiment. Without such replication, the claim is uncorroborated rather than contradicted. Expression compendia do not distinguish these cases because they catalogue datasets rather than claims^21^. Reanalysis of public raw data has recently questioned an entire class of published conclusions^22^, while editorials in this and other journals have emphasized replication and adequate statistical power as foundations of plant science^23–26^. What has been missing is a resource that measures corroboration itself.

Here we present PLEASE, the PLant Evidence Atlas for StrEss, a claim-centred atlas that makes independent support searchable through three design decisions. First, accessions, reanalyses and analytical routes derived from the same biological material are counted once as a study family, the unit of independent evidence. Second, a two-track ontology links source-located published claims to data-derived testable propositions and preserves the analytical provenance of every evidence record. Third, a portability framework asks whether independent context-matched evidence exists and, if so, whether it agrees, factorizing portability into match coverage and conditional agreement. PLEASE shows that a calibrated core of stress programmes survives dependency collapse, but corroboration falls sharply with contextual specificity as comparable independent experiments become scarce. Together, these measurements reveal where analytical depth is high but biological independence is low and translate those evidence gaps into priorities for validation, replication and future experiments.

## Results

### PLEASE separates biological independence from analytical depth

We searched eight public repositories and catalogues for plant stress transcriptomes with matched controls and froze the corpus on 27 August 2026 (Fig. 1a; Supplementary Fig. 1a; Supplementary Table 1). The three core sources, the Gene Expression Omnibus (GEO), the European Nucleotide Archive and Sequence Read Archive (ENA/SRA) and PubMed, yielded 26,575 candidate study clusters, of which only 2,151 were found by more than one source (Fig. 1b). Source complementarity was substantial, with the last source added contributing a median 35.9% of clusters across all possible orders (Supplementary Fig. 1b, c). PLEASE therefore integrates eight sources within a documented and reproducible search boundary.

**Fig. 1:**
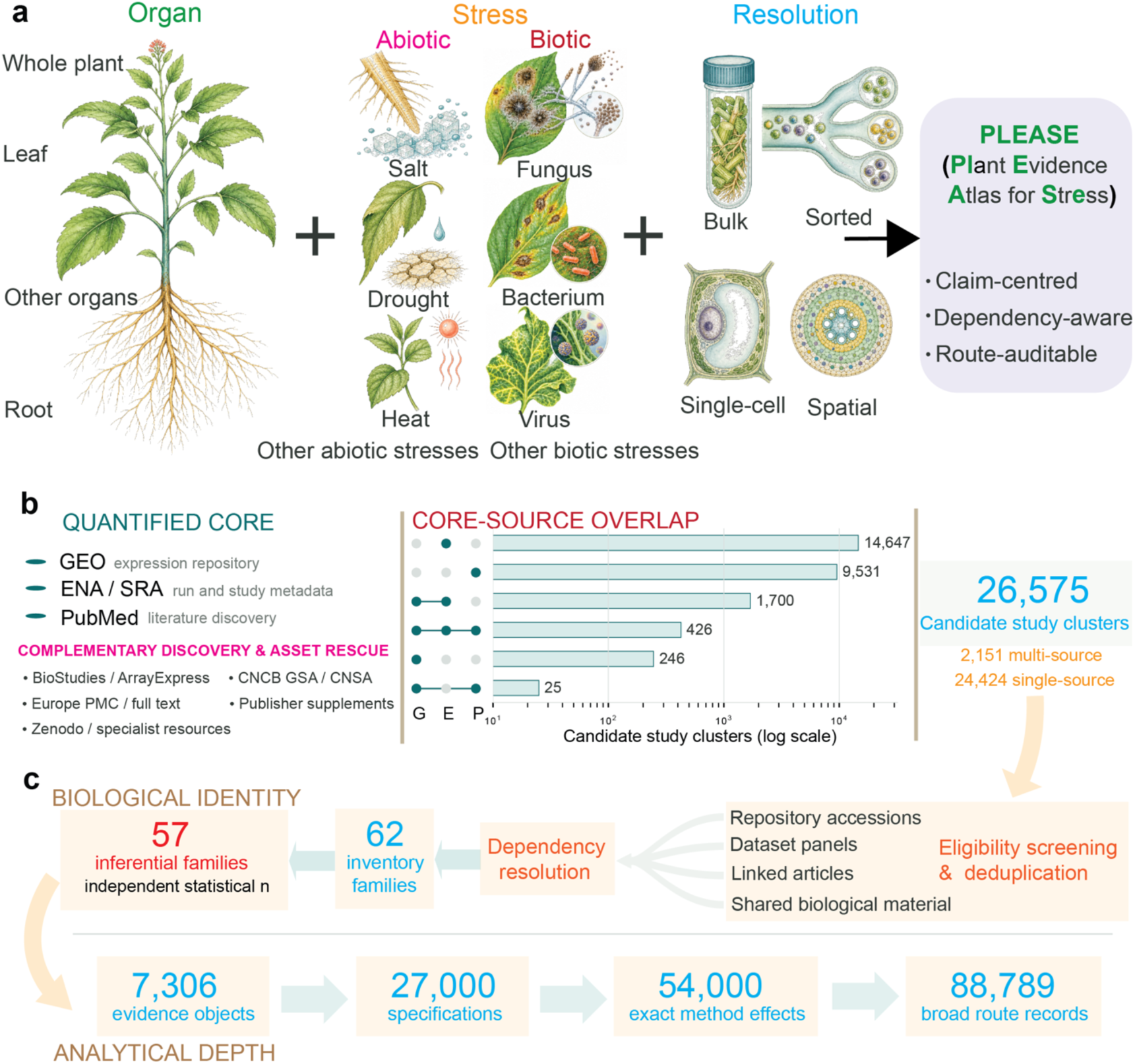
PLEASE separates biological independence from analytical depth. **a**, Biological scope of PLEASE. Public transcriptomic evidence spans plant organs, abiotic and biotic stresses, and bulk, sorted or laser-captured, single-cell or single-nucleus and spatial measurements, organized as a claim-centred, dependency-aware and route-auditable resource. **b**, Multi-source discovery. GEO, ENA/SRA and PubMed form the quantified search core. The UpSet plot shows the observed intersections among 26,575 candidate study clusters, including 2,151 detected by more than one source. Complementary repositories, full-text searches and supplementary materials support discovery and asset recovery outside this shared denominator. **c**, Dependency-aware resolution of biological identity and analytical depth. Candidate records were linked across accessions, dataset panels, publications and shared biological material and resolved into 62 study families, 57 of which, comprising 41 abiotic and 16 biotic families, provided the independent units for inference. From this base, 7,306 audited evidence records expand to 88,789 broad route-level records across analytical specifications. The hierarchy separates biological independence from analytical depth without inflating the independent denominator.

Screening and dependency resolution converted this fragmented record space into independent evidence units (Supplementary Fig. 1d–f). Accessions, dataset panels and publications derived from shared biological material were merged into 62 study families, 57 of which were eligible for primary inference, comprising 41 abiotic and 16 biotic families. The other five either used immune-activation proxies without a live pathogen or lacked biological replication and were retained for sensitivity or coverage analyses. These families span 17 species, 19 organs, 33 stress or treatment conditions and five measurement resolutions (Fig. 1a, c; Supplementary Fig. 1g; Supplementary Data 1).

PLEASE then links this independent evidence to two evidence tracks (Fig. 1c; Supplementary Table 2). The published track contains 71 harmonized claims, while the data track contains 1,811 biological coordinates (programme, stress, organ and cell lineage) and 2,543 directional propositions supported by 7,306 audited evidence records. These expand to 88,789 broad route-level records across analytical specifications, and 1,032 crosswalk links connect published claims to the propositions that can test them. The resulting hierarchy separates analytical depth from biological independence and allows each conclusion to be traced to both its analytical provenance and its independent experimental support.

### A calibrated core of programmes survives dependency collapse

We tested 20 response programmes defined before any atlas data were analysed, comprising 10 abiotic and 10 biotic programmes, each with 5–19 Arabidopsis genes projected into other species by orthology^27–30^. After collapsing dependent evidence to one vote per family and programme, recurrent and direction-consistent responses remained across the 57 independent families (Fig. 2a, b). Heat shock was evaluable in 35 of 41 abiotic families, with 82% of resolved families showing induction, while reactive oxygen species (ROS), abscisic acid (ABA) and osmotic programmes showed 76%, 75% and 81% directional agreement, respectively. All 10 biotic programmes were evaluable in all 16 biotic families, with agreement from 60% to 85%. Within-family route concordance was high, with medians of 0.81 for abiotic and 0.91 for biotic evidence (Fig. 2c, d).

**Fig. 2:**
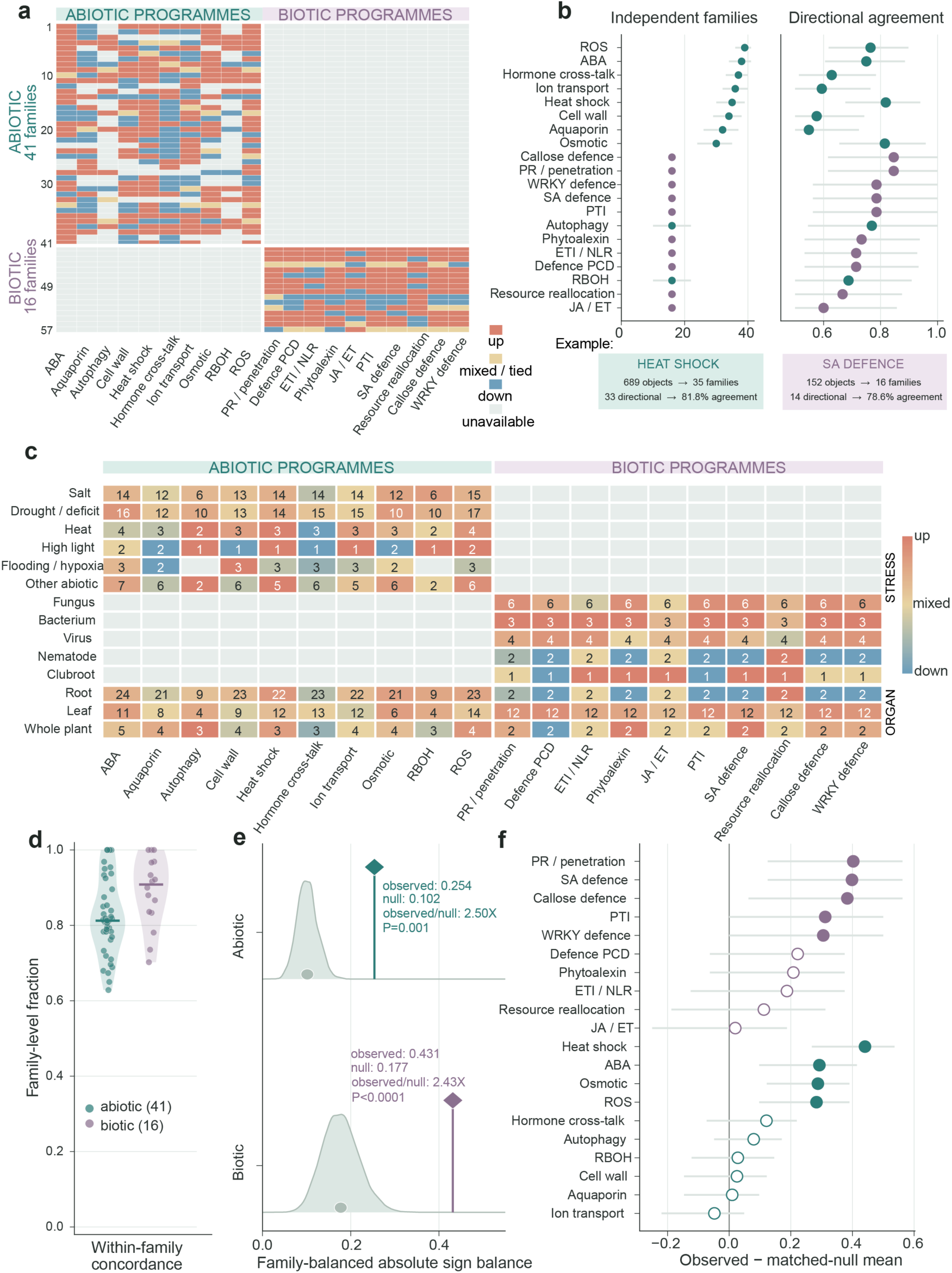
Independent families support a calibrated core of stress programmes. **a**, Directional map across 57 independent study families and 20 predefined response programmes. Dependent evidence was collapsed within each family so that every family contributes one equal-weight directional vote per evaluable programme; colours give the family-level direction (up or down), mixed or tied evidence, or unavailable combinations. Many unavailable cells are structural, reflecting the separate evaluation of abiotic and biotic programmes within their respective stress categories. **b**, Recurrence and directional agreement of the programmes after collapsing. Left, the number of independent families with evaluable evidence; right, agreement with the dominant direction among directional families, with 95% intervals from 5,000 family bootstraps. Heat shock illustrates the distinction between record abundance and independent support: 689 evidence records resolve to 35 families, of which 33 had a resolved direction, 81.8% of them were upward. **c**, Context-resolved programme architecture. Rows are stress and organ contexts, columns are programmes; numbers indicate contributing families and colour indicate the balance of family-level directions. **d**, Within-family analytical robustness. Each point is one family and summarizes directional concordance across analytical routes (median family-direction concordance 0.81 abiotic, 0.91 biotic). **e**, Observed family-balanced absolute sign balance (diamonds) against null distributions from expression- and precision-matched random gene sets (shaded densities; grey circles mark null means): abiotic 0.254 versus 0.102 (2.50-fold; one-sided *P* = 0.001; 1,000 draws), biotic 0.431 versus 0.177 (2.43-fold; *P* < 0.0001; 20,000 draws). **f**, Programme-level observed minus matched-null mean, with intervals giving the transformed central 95% null range. Filled symbols mark the nine programmes passing strict, coarse and expression-only matching.

This recurrence exceeded matched random gene sets (Supplementary Fig. 2). The family-balanced recurrence statistics were 0.254 versus a null mean of 0.102 for abiotic responses and 0.431 versus 0.177 for biotic responses, corresponding to 2.50-fold and 2.43-fold enrichment, respectively (*P* = 0.001 and *P* < 0.0001). The result persisted in every leave-one-family-out evaluation (Fig. 2e). Nine programmes passed all three matching schemes, including heat shock, ABA, osmotic and ROS responses, as well as pathogenesis-related, salicylic-acid, callose, pattern-triggered and WRKY-regulated defence programmes (Fig. 2f).

Programme directions were also stable in time and across the corpus. Predicted solely from families released earlier, they were correct in 0.62 of abiotic and 0.71 of biotic cases. Directions fixed from the first 13 primary biotic families were retained for nine of 10 programmes after five additional families were integrated, and removing any single species, organ or measurement resolution retained at least 78% of directions (Supplementary Fig. 3). The remaining question is how far this support persists as claims become more context specific.

### Corroboration collapses as claims gain specificity

For each target family, we asked whether at least two other families provided resolved directions at the same biological coordinate and whether the majority agreed with the target. Match coverage (*C*) measures the availability of independent, context-matched evidence, conditional accuracy (*A*) measures agreement given a match, and evidence portability (*P* = *C* × *A*) combines the two. Each target family contributes equally to these summaries. Across four nested levels of specificity, observed coordinates increased from 20 at the programme level to 249 after adding stress, 406 after adding organ and 1,023 after adding cell lineage (Fig. 3a).

**Fig. 3:**
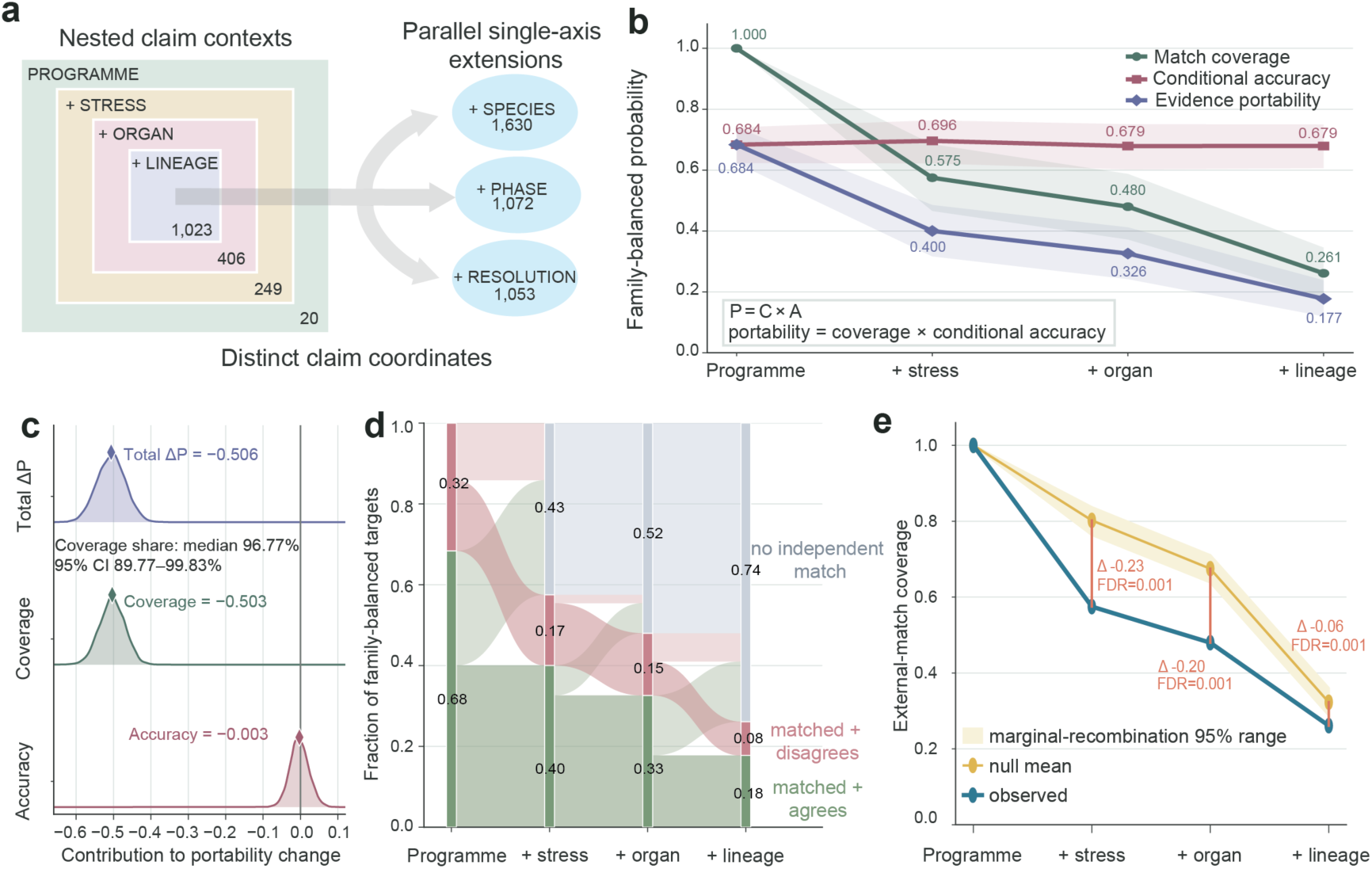
Independent coverage declines as claims become context-specific. **a**, Nested claim contexts. The primary series restricts each programme by stress, organ and lineage, expanding the claim space from 20 programme-level coordinates to 249, 406 and 1,023, respectively; species, phase and measurement resolution are parallel refinements (1,630, 1,072 and 1,053 coordinates, respectively) analysed as sensitivity branches. **b**, Match coverage *C*, conditional directional accuracy *A* and evidence portability *P* = *C* × *A* across the series. Coverage is the family-balanced probability that a target can be evaluated against at least two other independent context-matched families; accuracy is the probability of directional agreement given such a match. Shading, 95% intervals from 5,000 paired family bootstraps. **c**, Exact symmetric decomposition of the portability change from programme-level to lineage-specific claims. Densities show 5,000 family-bootstrap replicates, diamonds indicate point estimates and the vertical line marks zero. Coverage accounts for a median 96.8% (95% interval 89.8–99.8%) of the loss. **d**, Evidence-state composition. Family-balanced targets are matched with agreement, matched with disagreement or without an independent match; the unmatched fraction rises from 0 to 0.43, 0.52 and 0.74. **e**, Observed coverage (blue) against the marginal-recombination null (yellow line and shading indicating mean and 95% range, respectively), which preserves the frequency of every context label while disrupting their joint association. Observed coverage is lower at every rank (one-sided lower-tail P, Benjamini–Hochberg-adjusted across ranks, 0.001), and the coverage slope is steeper than under the null (*P* = 0.003); the portability slope tracks the null (*P* = 0.264) because matched directions hold up better than under it.

Coverage fell from 1.000 to 0.575, 0.480 and 0.261, whereas conditional accuracy remained at 0.684, 0.696, 0.679 and 0.679. Portability consequently fell from 0.684 to 0.177 (Fig. 3b). An exact symmetric decomposition assigned −0.503 of the −0.506 change to coverage and only −0.003 to accuracy, with coverage accounting for a median 96.8% of the loss across 5,000 family bootstraps (Fig. 3c). With increasing specificity, claims became unmatched rather than discordant. The unmatched fraction rose from 0 to 0.43, 0.52 and 0.74 (Fig. 3d). These states imply different scientific needs. An unmatched cell-lineage response requires new sampling. Disagreement among matched families may reflect biological heterogeneity, analytical sensitivity or annotation uncertainty and therefore requires further diagnosis.

Some coverage loss is expected as more contextual dimensions must match. We therefore compared observed coverage with a null model that preserves the frequency of each context label within programmes but recombines labels independently. Observed coverage was below expectation at every context rank, with 0.575 versus 0.802, 0.480 versus 0.675, and 0.261 versus 0.323 [false-discovery-rate (FDR)-adjusted *P* = 0.001 for each comparison]. The decline was steeper than expected (slope *P* = 0.003; Fig. 3e). The coverage deficit was robust to a tuple-preserving null and to stricter requirements of three or five training families per match (Supplementary Fig. 4). Combinatorial sparsity therefore explains only the baseline decline. The observed sampling structure further reduces independent corroboration beyond that expected from specificity alone.

### Sparse sampling sets the scale of the evidence gap

The coverage deficit reflects sparse independent sampling of biological contexts. The 57 families formed 170 family-context associations across 116 observed contexts defined by stress, organ, lineage and measurement resolution. Of these, 87 contexts (75.0%) contained a single family, 15 contained two and only 14 contained at least three, which is the minimum needed to evaluate a target against two independent families (Fig. 4a). Allocation concentration matched a degree-preserving null, with a Herfindahl–Hirschman index of 0.01671 versus 0.01665 and *P* = 0.436 (Fig. 4b). The scarcity therefore reflects both how few contexts each study covers and how rarely the same context is revisited independently.

**Fig. 4:**
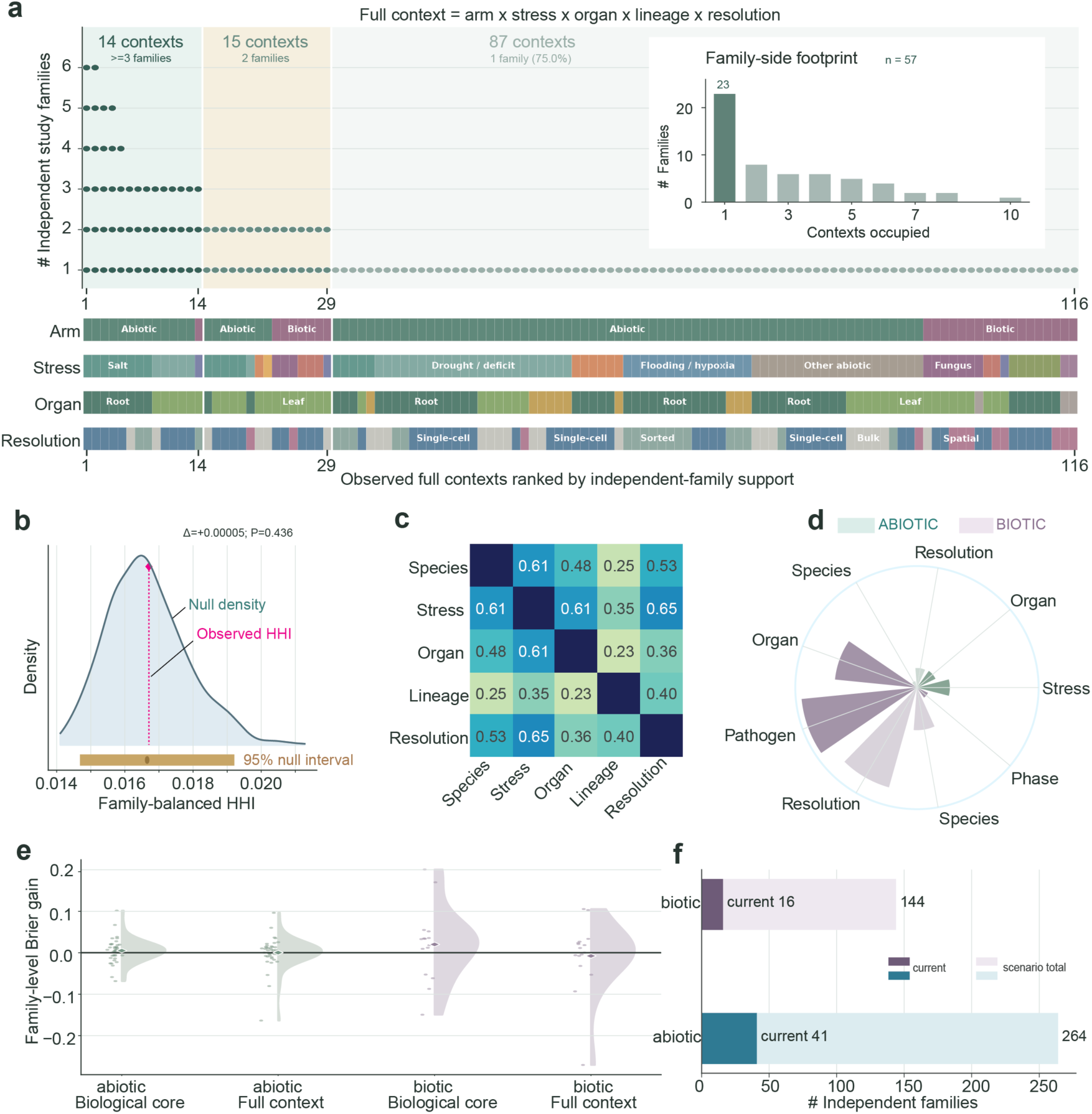
Sparse context occupancy structures the evidence and limits out-of-family prediction. **a**, Occupancy of the observed full-context space by 57 independent families. A full context is stress category, stress, organ, lineage and measurement resolution; the 57 families form 170 family–context incidences across 116 contexts, ranked by independent-family support, each stacked point being one family. Eighty-seven contexts (75.0%) hold a single family, 15 two and 14 three or more. Tracks show stress category, stress, organ and resolution; the inset gives contexts per family. **b**, Family-balanced allocation concentration (Herfindahl–Hirschman index) against a degree-preserving edge-swap null that fixes families per context and contexts per family: observed 0.0167 (dotted line) versus null mean 0.0166 (density; bar, central 95% interval; one-sided upper-tail *P* = 0.436). **c**, Pairwise Cramér’s V among species, stress, organ, lineage and resolution computed from the family–context incidence table; several axes are strongly coupled. **d**, Shapley decomposition of leave-one-family-out Brier-score reduction among context variables (sector radius, absolute contribution; colour, stress category; dark and pale, positive and negative). **e**, Prediction on wholly held-out families. Biological-core and full-context models are compared with a programme-only baseline (abiotic core, stress and organ; full, plus resolution and species scope; biotic core, organ and pathogen class; full, plus resolution, species scope and phase). Positive Brier gain favours added context; bars give 95% family-bootstrap intervals. **f**, Evidence-expansion scenarios. Resolving the observed gains at 80% power at the observed variance would require about 264 abiotic and 144 biotic families, compared with 41 and 16 available.

Context axes were also strongly coupled, with Cramér’s V of 0.61 to 0.65 for stress with organ, species and measurement resolution (Fig. 4c), making their individual contributions difficult to separate. We therefore tested whether context improved prediction of programme directions in wholly held-out families. Adding the biological core produced Brier-score gains of 0.0050 for abiotic and 0.0201 for biotic responses, both below the minimum detectable gains of 0.0127 and 0.0601, respectively, at the current sample sizes (Fig. 4d, e). At the observed gains and variance, achieving 80% power would require approximately 264 abiotic and 144 biotic families, compared with the current 41 and 16, corresponding to about sixfold and ninefold expansion, respectively (Fig. 4f). Sparse sampling therefore not only limits context-specific corroboration but also quantifies the independent evidence needed to resolve context effects.

### Cross-domain transfer separates shared direction from molecular support

We next tested whether evidence from one stress category could predict programme directions in the other. Family-balanced transfer accuracy was 0.759 at the global-programme rank, 0.750 at shared functional axes and 0.748 with organ or resolution matching, but fell to 0.615 at the cell–organ–lineage rank, where only 13 families were evaluable (Fig. 5a, b). Among the 13 families evaluable at all four ranks, accuracy declined by 0.068 per rank, with a 95% family-bootstrap interval of −0.142 to 0.010 and a rank-permutation *P* = 0.023 (Fig. 5c, d). Context matching did not improve accuracy over the source-category global-direction baseline, and source directions remained stable when individual stress domains were removed. The transferable signal was therefore more consistent with broad programme direction than with context-specific direction.

**Fig. 5:**
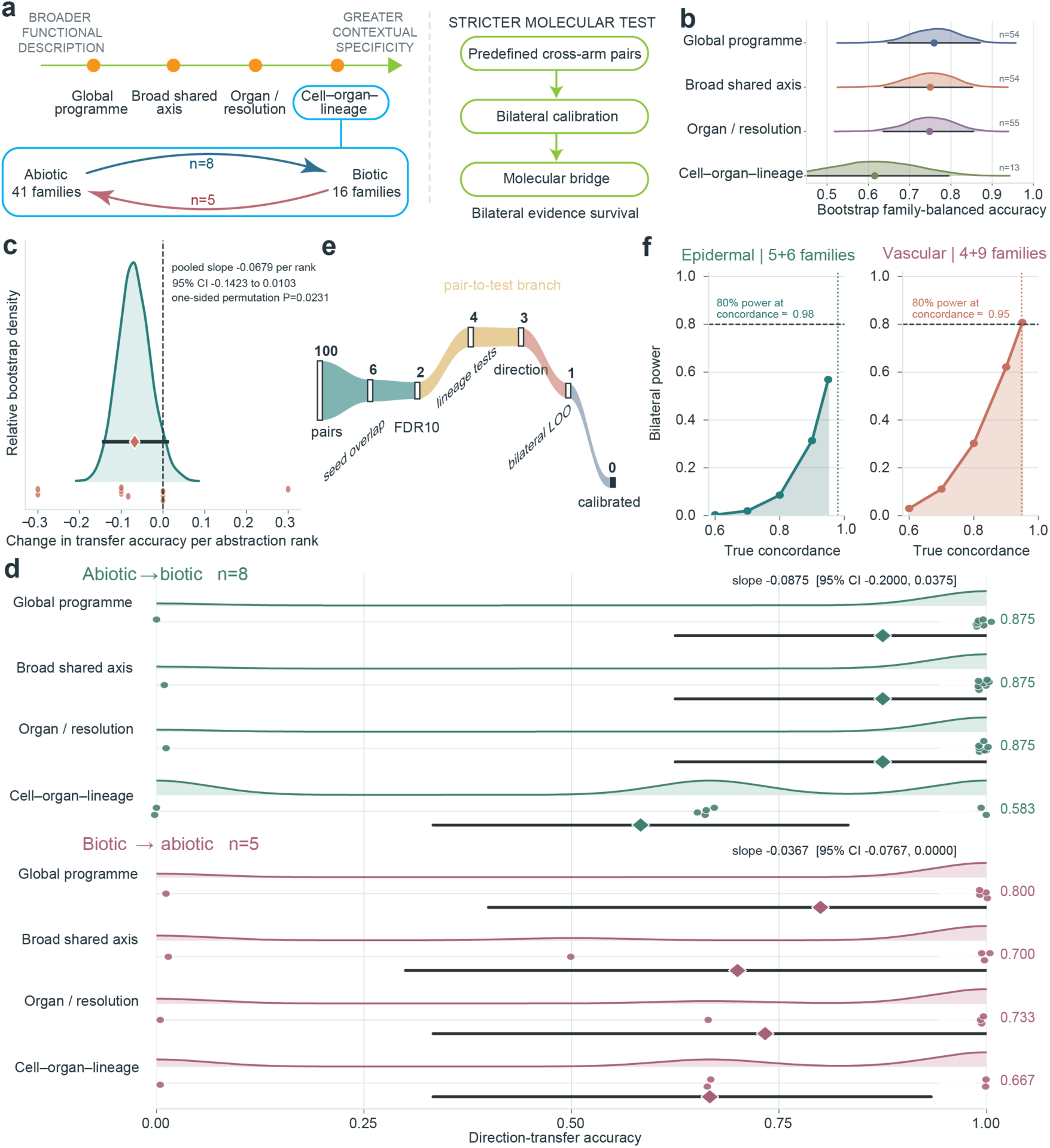
Cross-domain transfer and molecular calibration resolve different levels of support. **a**, Framework. Directional transfer between abiotic and biotic evidence is evaluated at four ranks of contextual specificity with target families held out in their entirety; 13 target families (eight biotic targets predicted from abiotic evidence and five abiotic targets predicted from biotic evidence) have the matched information required for gradient analysis. A separate, stricter analysis asks whether predefined cross-category programme pairs survive bilateral molecular calibration. **b**, Family-balanced direction-transfer accuracy at the four ranks (0.759, 0.750, 0.748 and 0.615); densities show 10,000 family-bootstrap distributions and numbers give evaluable target families. **c**, Matched-family gradient in the 13 families evaluable at every rank. The density shows the pooled slope over 10,000 family bootstraps; diamond and interval give the observed slope (−0.0679 per rank; 95% interval −0.1423 to 0.0103). A family-preserving permutation test gives one-sided *P* = 0.0231. **d**, Direction-specific transfer in the eight biotic and five abiotic matched targets; dots are family accuracies, curves are the bootstrap distributions of the family-balanced mean, diamonds and intervals are the mean and 95% interval, respectively; direction-specific slopes are printed on the panel. **e**, Survival of candidate molecular bridges: 100 predefined pairs, six with direct seed-gene overlap, two after 10% false-discovery control, four prespecified pair-by-lineage tests, three direction-concordant, one stable under bilateral leave-one-family-out analysis and zero passing bilateral calibration. Ribbon width tracks surviving units; the increase from two pairs to four tests is the predefined lineage branching, not additional support. **f**, Power for the bilateral directional criterion alone, which bounds full-test power from above, at the available family numbers: about 0.98 (epidermal) and 0.95 (vascular) true concordance is required for 80% power.

Shared direction did not translate directly into calibrated molecular support. Of 100 predefined abiotic × biotic programme pairs, six shared seed genes and two passed overlap FDR control, which are autophagy with defence-associated programmed cell death and the respiratory burst oxidase homologue (RBOH) programme with callose defence. Four predefined lineage-level tests yielded three direction-concordant results, one stable to family removal in both stress categories, but none passing bilateral matched-null calibration (Fig. 5e). With four to nine families per stress category, achieving 80% power for bilateral directional concordance required true concordance of 0.95–0.98 (Fig. 5f). Cross-domain evidence therefore supports broad programme direction, while specific shared molecular relationships require denser independent sampling.

### Evidence fingerprints define research priorities

PLEASE turns these evidence distinctions into six-component evidence fingerprints, summarizing independent corroboration, directional agreement, context breadth, analytical stability, annotation adequacy and evidence confidence (Fig. 6a, b). For example, the broad heat-shock programme contains 689 records from 35 families and provides a foundation for mechanistic work, whereas heat-shock induction in heat-stressed leaves contains 91 records from one rice experiment (GSE74793)^31^ and calls for independent replication (Fig. 6c). Salt-responsive root-hair ROS is directionally consistent across families but sensitive to analysis, making analytical re-evaluation a priority (Fig. 6d). Salicylic-acid defence is supported by 16 families overall, while its bacterium–leaf–mesophyll context rests on 48 records from two families and remains a scoped hypothesis for validation (Fig. 6e).

**Fig. 6:**
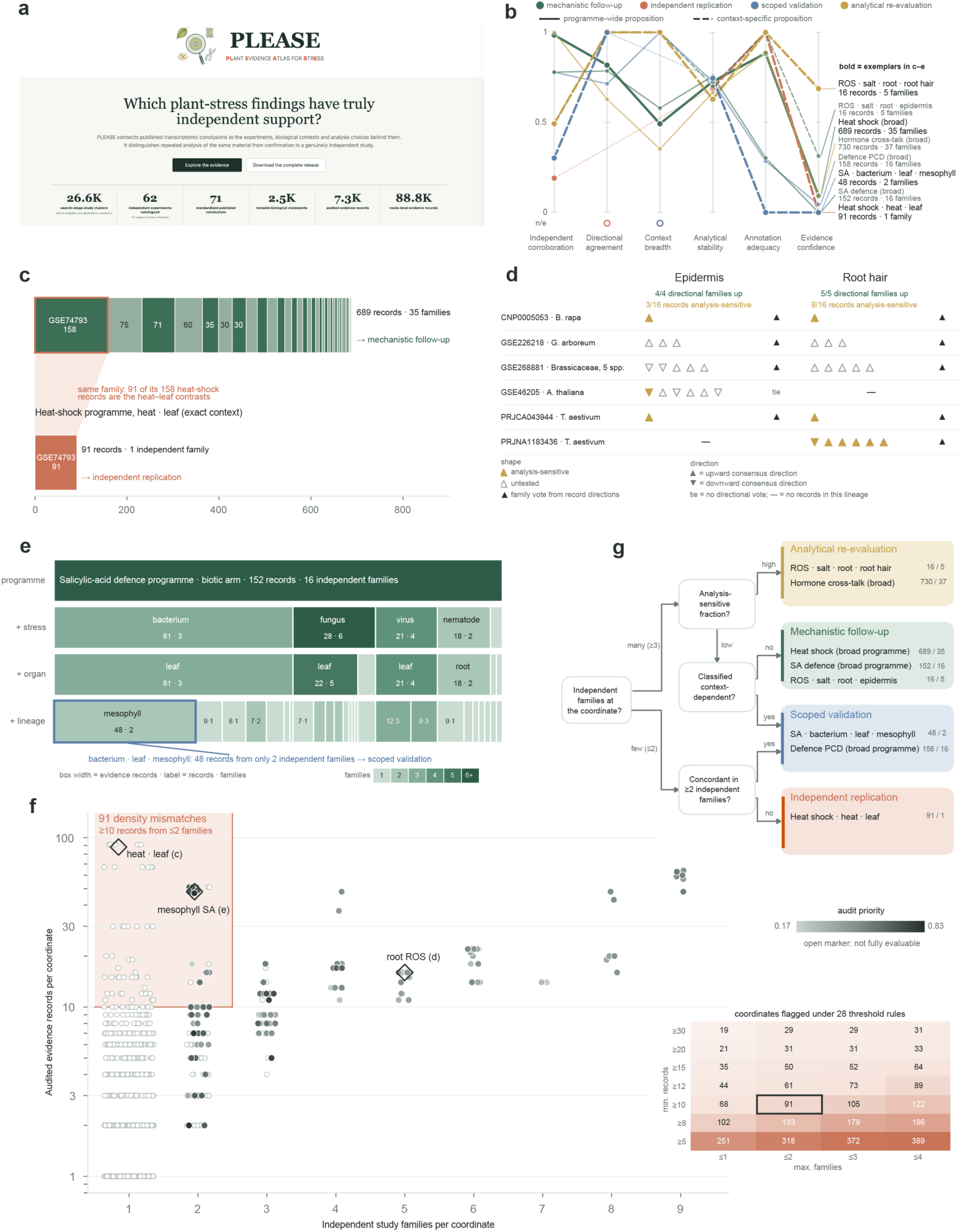
Evidence fingerprints translate corroboration deficits into distinct research priorities. **a**, Entry page of the PLEASE web interface. **b**, Evidence fingerprints of eight worked propositions across six components. The context-breadth axis measures consistency among evaluable records; unresolved records are excluded. It does not count sampled contexts. Lines are coloured by the action recommended in **g** (green, mechanistic follow-up; clay, independent replication; blue, targeted validation; gold, analytical re-evaluation); solid, programme-wide; dashed, context-specific; thick lines and bold labels, the exemplars in **c–e**. Open circles mark non-evaluable components (dotted segments bridge them); labels give records and families. **c**, Heat-shock evidence: 689 records from 35 families for the programme versus 91 records from one family (GSE74793) for the heat · leaf context; one bar segment per family, with the outlined segment marking the 91 records within that family’s 158 heat-shock records. **d**, Salt-responsive root ROS evidence in epidermis and root hair (16 records from five families per lineage, six families in total), one glyph per record grouped by family: upward- and downward-pointing triangles indicate upward and downward consensus direction, respectively; gold fill, analysis-sensitive records (3 of 16 and 8 of 16); open, untested. Black symbols give the family vote (a tie casts no vote); a dash marks a family without records in that lineage. **e**, Salicylic-acid defence evidence (152 records, 16 families) partitioned by stress, organ and lineage; box width is proportional to records and colour to families (1 to ≥6); the bacterium · leaf · mesophyll box (48 records, two families; outlined) is a scoped hypothesis. **f**, All 1,065 observed programme × stress × organ × lineage coordinates by independent families (jittered) and records (log scale). Filled markers, the 145 coordinates with a complete reanalysis-priority index (shaded 0.17–0.83); open markers, incomplete. The shaded region holds the 91 density mismatches (≥10 records, ≤2 families); diamonds mark the coordinates in **c–e**. Inset, coordinates flagged under 27 alternative rules and the prespecified rule (outlined). **g**, Decision tree from fingerprint components to the recommended next step; “many” families mean at least three, and leaves list the eight cases as records. Recommendations identify evidence-supported research priorities rather than verdicts on whether a proposition is true.

The published track connects these measurements directly to conclusions in the source literature (Supplementary Data 2). Of 73 assertions from 31 publications, 43 map to 878 atlas propositions and 18 specify a direction. The atlas reproduces 13, finds mixed evidence for four and opposing evidence for one. Ten are supported in the claimed direction by a second independent experiment, but none by a third. Replication debt extends the same distinction across the atlas. Among 1,065 observed coordinates, 91 meet the density-mismatch rule of at least 10 evidence records from no more than two families, and the 20 most indebted coordinates derive from a single family (Fig. 6f, Supplementary Fig. 5). The audit-priority index is available for 145 coordinates and correlates only moderately with replication debt at Spearman *ρ* = 0.43, showing that evidence concentration and methodological concern are distinct problems. A decision tree maps these patterns to mechanistic follow-up, scoped validation, independent replication and analytical re-evaluation (Fig. 6g).

PLEASE also registers its own predictions. A total of 1,599 programme-direction predictions at four specificity ranks were fixed on 27 August 2026 for evaluation against studies released later. The web resource links searchable propositions, published claims, independent studies, replication gaps and downloadable evidence tables (Supplementary Fig. 6). PLEASE therefore couples a map of current evidence with a defined route to improving it.

## Discussion

PLEASE establishes a new kind of resource for plant biology, an atlas of evidence rather than expression. Its unit is the claim and its denominator is the independent experiment. By separating analytical depth from biological independence, PLEASE changes how apparently well-supported findings are interpreted. For example, broad heat-shock responses are supported across many independent experiments and can motivate mechanistic follow-up, whereas heat-shock induction in heat-stressed leaves rests on one experiment and calls for replication. Salt-responsive root-hair ROS is supported across families but is sensitive to analytical choices, pointing instead to re-evaluation. Together, these examples show that similar volumes of transcriptomic evidence can imply very different levels of biological corroboration.

Using the independent experiment as the denominator reveals a calibrated but bounded core of plant stress biology. Predefined response programmes remain recurrent after dependent evidence is collapsed, their directions are stable across time and corpus perturbations, and broad programme direction transfers between abiotic and biotic evidence. The transferable signal is therefore strongest at the programme level. More specific conclusions require denser independent sampling.

The central finding is a mismatch between the resolution of biological claims and that of independent sampling. As claims specify stress, organ and cell lineage, independent match coverage falls sharply while directional agreement remains stable. Three quarters of sampled contexts contain only one study family, and observed coverage falls below that expected from context frequencies alone. The main limit on context specific corroboration is therefore not pervasive disagreement between experiments, but the absence of comparable independent experiments. The curated published claims show the same pattern, with no tested directional assertion supported by more than two independent experiments. The literature has accumulated far more analytical repetition than independent corroboration at the resolution at which many conclusions are now stated.

PLEASE turns this mismatch into distinct research priorities. Evidence fingerprints distinguish lack of replication from analytical sensitivity, restricted context breadth and directional disagreement, while replication debt identifies conclusions for which many analytical records still derive from few independent experiments. These states require different responses. Reanalysis can resolve analytical sensitivity, sampling a new context broadens biological coverage, and independently revisiting an existing context strengthens corroboration. By separating these contributions, PLEASE shows where another analysis is informative, where another context is needed and where only an independent experiment can close the evidence gap.

PLEASE also makes this evidence framework testable as literature grows. New studies can enter under the same evidence model^32^, while the registered predictions provide prospective tests under fixed rules. More broadly, any corpus in which public data are reanalysed more often than experiments are repeated can be examined using the same three elements, i.e. an independence denominator, a claim-centred ontology and an exactly factorized portability metric. For plant stress biology, PLEASE therefore does more than organize existing transcriptomes. It defines which conclusions rest on independent evidence, exposes where apparent depth is analytical rather than biological, and shows where new experiments can most increase what the field can claim with confidence.

## Methods

### Search sources and candidate discovery

Eight public repositories and catalogues were searched for plant stress transcriptomes with a matched control. GEO^6^, ENA/SRA^7^ and PubMed were treated as core sources because their records could be linked through a common retrieval unit. BioStudies/ArrayExpress^33^, CNCB GSA and CNSA/CNGB^34^, scPlantDB^9^ and PlantscRNAdb^10^ were used as complementary sources. Europe PMC full text^35^, publisher supplements, a curated Zenodo collection of processed single-cell objects and the PsctH catalogue supported discovery and asset recovery. Queries combined plant taxon terms with abiotic and biotic stress terms.

Core-source records were linked by exact identifiers into candidate study clusters. Cluster incidence across the three core sources was used to calculate multi-source and single-source counts, source-order accumulation curves and the terminal-source share. Lincoln–Petersen and Chao2 estimators^36^ were also applied to the incidence matrix as capture–recapture diagnostics. Because repository cross-listing violates independent capture and the two estimators gave strongly divergent totals, corpus completeness was not inferred from these estimates. The corpus was therefore defined by its documented search boundary. Per-source search and screening counts are provided in Supplementary Table 1.

### Screening and eligibility

Candidate accessions were screened in four stages. Invalid identifiers were removed, aliases and cross-accession links were merged, and the remaining components were evaluated against predefined eligibility criteria. A component was eligible for inference when it provided an analysis-ready processed matrix or reconstructable counts, a verifiable stress-control design with resolvable sample metadata, at least two biological replicates per group or valid independent samples for cell-resolved data, and a plant stress within scope. Duplicate or dependent records were not counted separately and were merged into the corresponding study family.

In the abiotic arm, 1,952 accession records resolved to 1,642 canonical components after identifier checking and merging. Forty-one components were admitted for inference, while GSE121619 was retained as a descriptive record without biological-replicate inference. A further 1,573 metadata-only records were retained in the public queue for future assessment (Supplementary Methods). The biotic arm was screened under the same criteria and comprised 21 study families, of which 16 were eligible for primary inference and five were retained for sensitivity or coverage analyses. These five families either used immune-activation proxies without a live pathogen or lacked biological replication.

### Dependency resolution into study families

Accessions, dataset panels, publications and reanalyses were merged into a single study family when they derived from the same biological material, sample collection or experimental data, as established from repository metadata and associated publications. Similar treatments, shared species or a common laboratory were not sufficient for merging, and multiple accessions were not treated as independent when they represented the same underlying experiment.

The 87 accessions in the corpus resolved to 62 study families, 15 of which contained two or more accessions. Of these, 57 families were eligible for primary inference, including 41 abiotic and 16 biotic families (Supplementary Data 1). The registry spans 17 species, 19 organ labels, 33 harmonized stress or treatment conditions, and five measurement resolutions, comprising bulk tissue or organ, sorted or enriched populations, laser-captured cell layers, single cells or nuclei, and spatial regions. The study family is the independent unit for all bootstraps, null models, held-out evaluations and *P* values.

### Curation of published claims

For each study family, associated publications were identified from repository links and full text. An assertion was admitted only when the authors explicitly interpreted a response to the focal stress or infection. Each assertion retained a source locator, a short anchor phrase and controlled values for entity, programme, direction, stress, organ, cell type or zone, time point, resolution, evidence mode and causal language (Supplementary Methods).

The published track contains 73 assertions from 31 publications associated with 30 abiotic inference families and the descriptive record GSE121619. Assertions were extracted through human reading, verified against the source text and adjudicated by a human curator. They were harmonized into 71 claims and linked to compatible atlas propositions through 1,032 claim-proposition pairs. Each mapped claim was connected to evidence from its source study and to compatible programme-level propositions. Directional comparisons refer to the mapped programme endpoint and do not retest narrower gene, metabolite or phenotype endpoints. Independent support was defined as the largest number of families supporting any direction-compatible linked proposition (Supplementary Data 2).

The curation protocol was independently benchmarked on checksummed full texts from 10 additional atlas families without web access. Their outputs were compared using a prespecified pairing score (Supplementary Table 3). Agreement was high for programme, direction and organ but lower for finer descriptors and for the set of propositions recovered. Model output was therefore used only to propose candidate annotations, and every registry entry was verified and adjudicated by a human curator. Controlled vocabularies followed community standards for plant anatomy, traits and evidence codes^37,38^.

### Response programmes

Twenty response programmes were defined before any atlas data were analysed. Ten abiotic programmes comprised heat shock, ABA, osmotic, ROS, hormone cross-talk, autophagy, RBOH, cell wall, aquaporin and ion transport, with 5–19 genes per programme. Ten biotic programmes comprised pathogenesis-related or penetration responses, salicylic-acid defence, callose defence, pattern-triggered immunity, WRKY regulation, defence-associated programmed cell death, effector-triggered immunity or nucleotide-binding leucine-rich repeat receptor signalling, indolic phytoalexin synthesis, sugar-transport resource reallocation and jasmonate or ethylene defence, with 9–16 genes per programme. Programme membership was assembled from canonical Arabidopsis pathway genes^27–30^.

A route contributed to a programme when at least three member genes were tested. Programme genes were transferred across species using within-species identifiers or predefined native sets, Ensembl Plants or curated orthologue sets, protein-sequence reciprocal best hits or family-level sequence similarity, and published cross-species hits where available^39,40^. The projection route was recorded for every effect. Four additional root-hair programmes were retained for resource queries, the resource-level replication-debt scan and the registered predictions. They were excluded from the primary recurrence, portability and prediction analyses.

### Analytical routes and evidence records

Each family was analysed using contrast-specific routes appropriate to its assay and data type. Count data, including replicate-aware pseudobulk profiles summed per biological sample and cell type^16,41,42^, were modelled using edgeR quasi-likelihood negative-binomial fits^43^ or limma on log counts per million after trimmed-mean-of-M-values (TMM) normalization, with or without voom precision weights^44,45^. Normalized-expression, microarray and author-transformed data were analysed using limma trend models. All comparison groups with sample-level data contained at least two biological replicates. For cell-resolved data, cell identities were evaluated through up to four annotation routes comprising source annotation, literature markers, three-route consensus and reconstructed author signatures, using all quality-controlled cells or balanced downsamples^46^.

Programme responses were evaluated with the competitive camera test and the self-contained fry test^47,48^. These paired tests were applied to 26,940 analytical specifications. A further 60 specifications derived from published summary statistics were evaluated using rank and signed-enrichment tests on author tables and were labelled separately.

An analytical specification was defined as one combination of normalization, annotation route and gene-set projection. The 27,000 specifications yielded 54,000 method-level effects, with additional sensitivity routes contributing to 88,789 broad route-level records. An audited evidence record represented one contrast, context and programme combination within a study family. Each of the 7,306 evidence records retained the consensus direction across its specifications, a strict significance gate requiring both gene-set tests to be significant in the same direction, and flags for direction and significance sensitivity across specifications.

### Family-level directions and recurrence calibration

Within each study family, directions were collapsed sequentially from method-level effects to analytical specifications, evidence records and study nodes. A study node represented one directional call for a dataset panel and contrast group at a biological coordinate. Study-node directions were then summed within each family and programme, and the sign of that sum defined the family vote. A zero sum was treated as a tie and contributed no direction. Dominant-direction agreement was calculated as the majority fraction among families with a resolved directional vote, with 95% intervals obtained from 5,000 family bootstraps^49^.

Programme recurrence was quantified as the mean absolute family-balanced sign balance across programmes. Unavailable and tied family-programme combinations therefore reduced the recurrence statistics, so the measure reflected both evaluability and directional consistency. Calibration used random gene sets matched to each programme within every analytical route for set size, tested universe, average expression and moderated precision. Three matching schemes were used, comprising strict 5 × 5 expression-precision strata, coarser strata and abundance-only matching. Random sets were processed through the same analytical tests and collapsing hierarchy as the predefined programmes. Empirical upper-tail *P* values were calculated from 1,000 abiotic and 20,000 biotic draws using the plus-one correction^50^. Programme-level false-discovery rates were controlled with the Benjamini-Hochberg procedure^51^, and robustness was assessed by leave-one-family-out analysis.

### Validation of programme directions

Four complementary designs tested whether programme directions generalized beyond the families used to define them. First, directions fixed from the 13 primary families among the first 14 integrated biotic families were evaluated in the three primary and two sensitivity-only families integrated subsequently. Second, leave-one-group-out analyses removed each species, organ or measurement resolution in turn, as well as all Arabidopsis families, and recomputed programme directions from the remaining families. Third, three abiotic studies with primary endpoints specified before integration provided 11 independent endpoint tests. Fourth, prequential evaluation ordered families by repository release date and predicted each family only from families released earlier, requiring at least three training families. Performance was compared with the family-balanced arm-wide direction baseline, with fixed and rolling cut-offs examined as sensitivity analyses.

In addition, 1,599 programme-direction predictions at four specificity ranks across all 24 resource programmes were fixed on 27 August 2026. Of these, 204 had a resolved direction supported by at least three training families. The accompanying protocol specifies eligibility of future studies, the primary endpoint and the confirmatory rule, which requires at least 10 eligible post-registration families including three per arm and a lower 95% family-bootstrap bound above 0.5.

### Independent matching and evidence portability

Within each stress category, propositions were evaluated at four nested specificity ranks comprising programme, programme plus stress, programme plus stress and organ, and programme plus stress, organ and lineage. For each target family, the reference set contained all other families with a resolved non-tied vote at the same biological coordinate. An independent match required at least two such reference families and a non-tied majority direction. The target family was always excluded from its own reference set.

Match coverage *C* was defined as the family-balanced fraction of target coordinates with an independent match. Conditional accuracy *A* was the family-balanced fraction of matched target coordinates whose direction agreed with the reference majority. Evidence portability *P* was the family-balanced fraction of all target coordinates that were both matched and correctly predicted. By construction *P* = *C* × *A*, all summaries gave equal weight to study families rather than analytical records.

The change in portability from programme level to lineage level was decomposed symmetrically into contributions from coverage and conditional accuracy, with the interaction divided equally between the two components using the two-component Shapley allocation^52^. Uncertainty was estimated from 5,000 paired family bootstraps of the family-by-rank quantities. Species, phase and measurement resolution were analysed as parallel refinements of the lineage rank rather than as additional ranks.

### Context-sparsity null models and direction baselines

The primary null permuted stress, organ, lineage and measurement-resolution labels independently within each stress category and programme stratum. This preserved the marginal frequency of every context label and each family’s node density while removing the observed joint context structure. Independent matching and prediction were then recomputed for each of 1,000 null iterations.

Observed coverage at each specificity rank was compared with its null distribution using one-sided lower-tail empirical *P* values. *P* values across ranks were adjusted using the Benjamini-Hochberg procedure, and the observed coverage slope across the four ranks was compared with the corresponding null slopes.

A tuple-preserving null served as a sensitivity analysis by shuffling complete context tuples rather than their component labels. The quantities preserved by the two null models are listed in Supplementary Table 4. Always-up, stress-category-majority, programme-majority and composition-expected direction baselines were evaluated on the same match mask with target families excluded. Sensitivity analyses repeated the portability calculations across all 24 resource programmes and with independent-match requirements of at least three or five training families.

### Sampling geometry

A full context was defined by stress category (abiotic or biotic), stress, organ, lineage and measurement resolution. Each study family was linked once to every full context that it occupied, producing a binary family by context incidence matrix with 170 incidences across 116 observed contexts. To prevent families occupying many contexts from contributing more total weight, each family was assigned unit mass divided equally among its occupied contexts. Context concentration was summarized using the Herfindahl Hirschman index calculated from these context masses.

A degree preserving null randomized the incidence matrix by double edge swaps within each stress category while preserving every family degree, context degree and the observed set of contexts fixed^53,54^. One thousand randomized matrices provided the null mean, central 95% interval and one-sided upper tail empirical *P* value for context concentration. Dependence among species, stress, organ, lineage and measurement resolution was quantified using Cramér’s V on the 170 family context incidences. A sensitivity analysis repeated these calculations after weighing each family’s incidences to unit mass.

### Context-informed prediction of programme directions

The prediction target was whether each family-programme vote agreed with the leave-family-out programme majority. L2-regularized logistic models^55^ were fitted at three levels of contextual information. The baseline model used programme identity alone. The biological-core model added stress and organ for abiotic evidence, and organ and pathogen class for biotic evidence^56^. The full model additionally included measurement resolution and species scope, with phase also included for biotic evidence.

All rows from each target family were held out together. Within the remaining training data, the inverse regularization penalty from 0.01 to 4 was selected using family-held-out tuning. Categorical predictors were one-hot encoded, with unseen categories ignored at prediction. Model performance was measured using family-balanced Brier scores^57^. Gains over the programme-only model were evaluated by resampling families 10,000 times.

An exploratory Shapley decomposition over moderator subsets was performed using fixed-penalty models. Minimal detectable Brier-score gains and evidence-expansion requirements were estimated using a normal approximation based on the observed family-level gain variance, 80% power and two-sided *α* = 0.05, with the observed gain used as the target effect (Supplementary Methods).

### Cross-domain transfer and molecular bridges

Directional transfer was evaluated between the abiotic and biotic stress categories at four ranks of contextual specificity comprising global programme, broad shared axis, organ or resolution matched axes, and cell organ lineage contexts. Programme directions were estimated from one stress category and evaluated in whole target families from the other, with each target family excluded from the source evidence. Family balanced accuracy was calculated at each rank and resampled 10,000 times over evaluable families.

The matched gradient analysis used the 13 target families evaluable at all four ranks. A least squares slope across ranks was calculated for each family, and the mean slope was evaluated using 10,000 family bootstraps and a 10,000 iteration within family rank permutation test. Source directions were also recomputed after removing each stress domain in turn to assess their stability. Molecular bridges were evaluated separately using 100 predefined abiotic and biotic programme pairs. Direct seed gene overlap was tested using a hypergeometric test against 27,655 *Arabidopsis thaliana* protein coding genes, with false discovery rate controlled at 10%. Programme pairs passing this screen were then evaluated for lineage specific directional concordance, bilateral leave one family out stability and bilateral calibration against the matched null. Power for the directional criterion was calculated from the available family numbers in each stress category using a binomial model and combined across the two categories. Because overlap, stability and programme calibration were additional requirements, this directional calculation provides an upper bound on power for the complete molecular test.

### Evidence fingerprints, replication debt and audit priority

Each evidence record carries five stored classifications. Replication status was classified as sufficient when at least two other families provided directional votes at the same coordinate and as limited otherwise. Context dependence was classified as detected when those families agreed in fewer than 75% of cases and not detected when agreement was at least 75%. Analytical sensitivity was classified as sensitive when direction or significance changed across analytical specifications, stable when at least two specifications agreed and passed the strict gate, untested when only one specification was available, and partially tested otherwise. Annotation support reflected the confidence of cell type mapping and was not applicable to records without cell type evidence. Evidence confidence was classified as high when replication was sufficient, agreement and the consensus direction fraction were at least 0.75, and the record was null calibrated, not power limited and analytically stable. Confidence was classified as moderate when replication was sufficient, agreement and the direction fraction were at least two-thirds, and the record was not untested. All other records were classified as limited.

For each proposition, the six component evidence fingerprint summarized independent corroboration as the log scaled family count, directional agreement among directional families, context breadth as the fraction of classified records without detected context dependence, analytical stability as the mean stability score, annotation adequacy as the fraction of records with high support or not applicable annotation, and evidence confidence as the fraction of records classified as moderate or high. Records with unresolved context dependence were excluded from the context breadth denominator.

For each observed stress category × stress × organ × lineage × programme coordinate with evidence records (*O*) and independent families (*F*), replication debt (*D*) was defined as

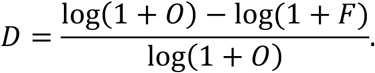

The prespecified density mismatch rule flagged coordinates with *O* ≥ 10 and *F* ≤ 2 . Twenty-seven alternative thresholds with *O* ≥ 5 to 30 and *F* ≤ 1 to 4 were evaluated as sensitivity analyses. The audit priority index was the mean of analytical sensitivity, directional discordance and annotation uncertainty, where directional discordance was one minus the majority fraction among at least two directional families. The index was calculated only when all three components were available. These quantities were used to distinguish mechanistic follow up, scoped validation, independent replication and analytical reevaluation.

### Web resource

PLEASE is a static web application with six views. These comprise an entry page with headline denominators, an evidence browser filtered by stress type, stress, organ, lineage, programme and direction, a published evidence view linking each harmonized claim to compatible propositions and supporting independent studies, a study registry with official repository links, a replication gap view ranking coordinates by replication debt, and a Methods and Downloads page providing analysis layers.

On the website, study families are labelled “independent experiments”, harmonized claims are labelled “standardized published conclusions”, propositions are labelled “testable biological statements”, the replication debt index is shown as the “replication gap score”, and the audit priority index is shown as “reanalysis priority”. All 62 study families are linked to their corresponding public repository records. Original expression and sequence data are not redistributed.

Releases are versioned, with this manuscript describing release 1.4.0. All counts displayed on the website are computed directly from the released tables, and new studies are admitted under the published eligibility protocol.

### Statistics and reproducibility

The study family was the independent sampling unit throughout, and prediction analyses held out entire families. Unless stated otherwise, confidence intervals were percentile intervals from family bootstraps. Empirical *P* values were one sided and used the plus one correction. Upper tail tests were used for recurrence and concentration, whereas lower tail tests were used for coverage depletion and the matched cross domain slope. Null reference intervals were distinguished from confidence intervals. Iteration counts for individual analyses are listed in Supplementary Table 5. Analyses were performed using Python 3.12 with pandas 3.0.5, NumPy 2.5.2, SciPy 1.18.1, scikit learn 1.9.0 and statsmodels 0.14.6, and R 4.4.3 with limma 3.62.1 and edgeR 4.4.0.

## Data availability

All data underlying the analyses in this study are freely available through the PLEASE resource at https://ai4lifesciences.github.io/PLEASE/. The resource provides the dependency-resolved study registry, harmonized published claims and atlas propositions, analytical evidence tables, programme definitions, analysis outputs and registered predictions, together with links to the original public repositories for all source studies. Original expression and sequence data are not redistributed.

## Code availability

All code used to build the resource and reproduce the analyses is freely available through the PLEASE resource at https://ai4lifesciences.github.io/PLEASE/. The resource provides the analysis code and associated release files required to reproduce the reported results. The resource is freely accessible without registration and will be maintained for the foreseeable future.

## Acknowledgements

The authors acknowledge the University of Sydney Research Data Store (RDS) and research computing infrastructure for providing data storage and computational resources used in this study. The authors also acknowledge Dr. Nathaniel Butterworth of Google Australia Pty Ltd for his help and valuable discussions.

## Author contributions

W.D. conceived the study and developed the methodology. W.D., Y.D. and P.Z. all contributed to the conception of the work. W.D. performed the analyses and wrote the initial manuscript. W.D., Y.D. and P.Z. interpreted the results and revised the manuscript.

## Funding

This work was supported by Google Australia’s Digital Future Initiative.

## Competing interests

The authors declare no competing interests.

## Notes

### Competing Interest Statement

The authors have declared no competing interest.

